## Supplemental Figures S1-S6 &Tables S1-S3 for "Depletion of aneuploid cells is shaped by cell-to-cell interactions"

Supplementary figures and tables

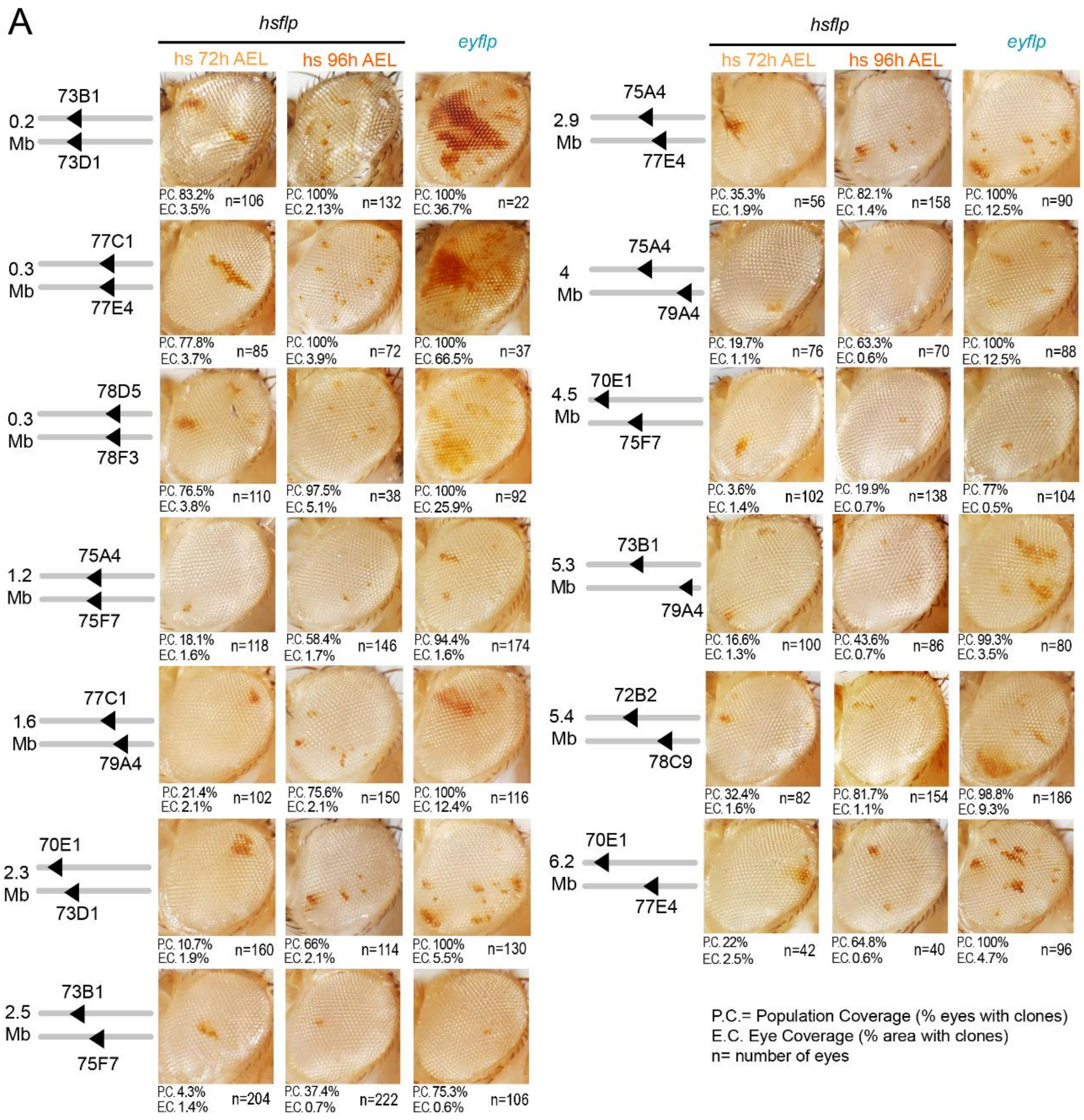


**Figure S1. Recombination efficiency relies on genomic distance** (related to Figure 1)

**(A**) Adult eyes with clones of cells labeled in red as a result of a recombination event between two RS-FRTs placed at the indicated genomic locations. Population and eye coverages (P.C. and E.C., respectively) and number of eyes scored for each FRT combination are shown.

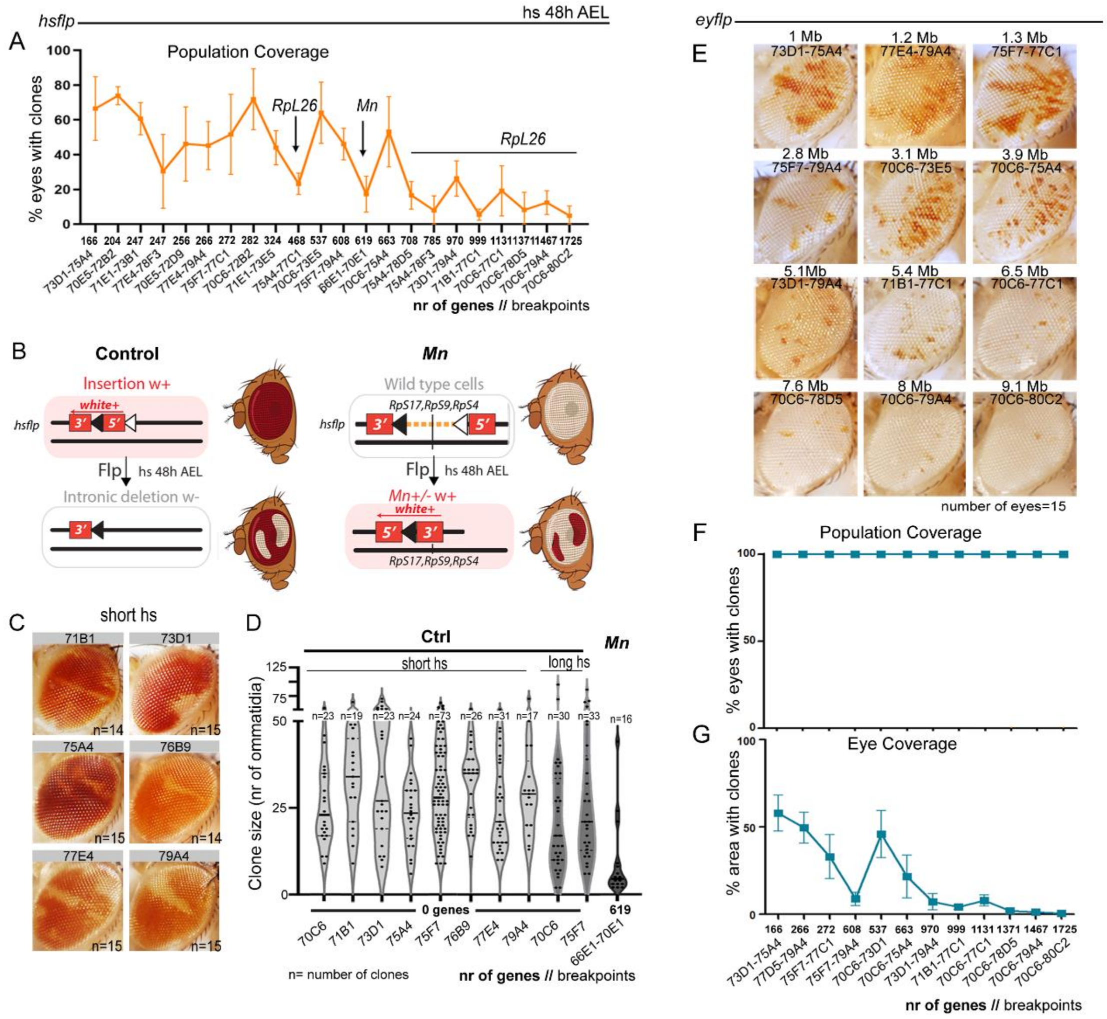


**Figure S2. The effect of genomic distance on recombination efficiency** (related to Figure 2)

(**A**, **F**, **G**) Plots representing the impact of the genomic distance (in number of genes) between the two FRTs on population coverage (% of eyes with clones, **A**, **F**) and eye coverage (% area with clones, **G**) of clones induced either acutely at 48h AEL (in orange, **A**) or chronically with the ey-FLP construct (in blue, **F**, **G**). Genomic location of each FRT combination is also indicated. (**B**) Drawings of clones of cells labeled in white (left) or red (right) as a result of a recombination event between two RS-FRTs located in the same chromosome (in *cis*). Clones acting as controls delete the 5’ intron of the *white* gene without affecting any gene (left), whereas clones inducing segmental monosomies and including *RpS17*, *RpS9* and *RpS4* (right) reconstitute the *white* gene. Recombination events are induced in founder cells during the growing period of the eye primordium and these cells proliferate and grow to give rise to clones visible in the adult eye. (**C**) Adult eyes with clones of cells labeled in white as a result of a recombination event within RS-FRTs placed at the indicated genomic locations. Clones in **A**, **C** and **D** were induced acutely at 48 h after egg laying (AEL). (**D**) Plot representing the size of control clones (in number of ommatidia) induced with the indicated RS-FRTs with short (light gray) and long (dark grey) heat-shock protocols (see Materials and Methods). Genomic location of RS-FRTs is indicated. Mean and SD are shown in **A**, **F**, **G**. Median is shown as a black line in **D**.

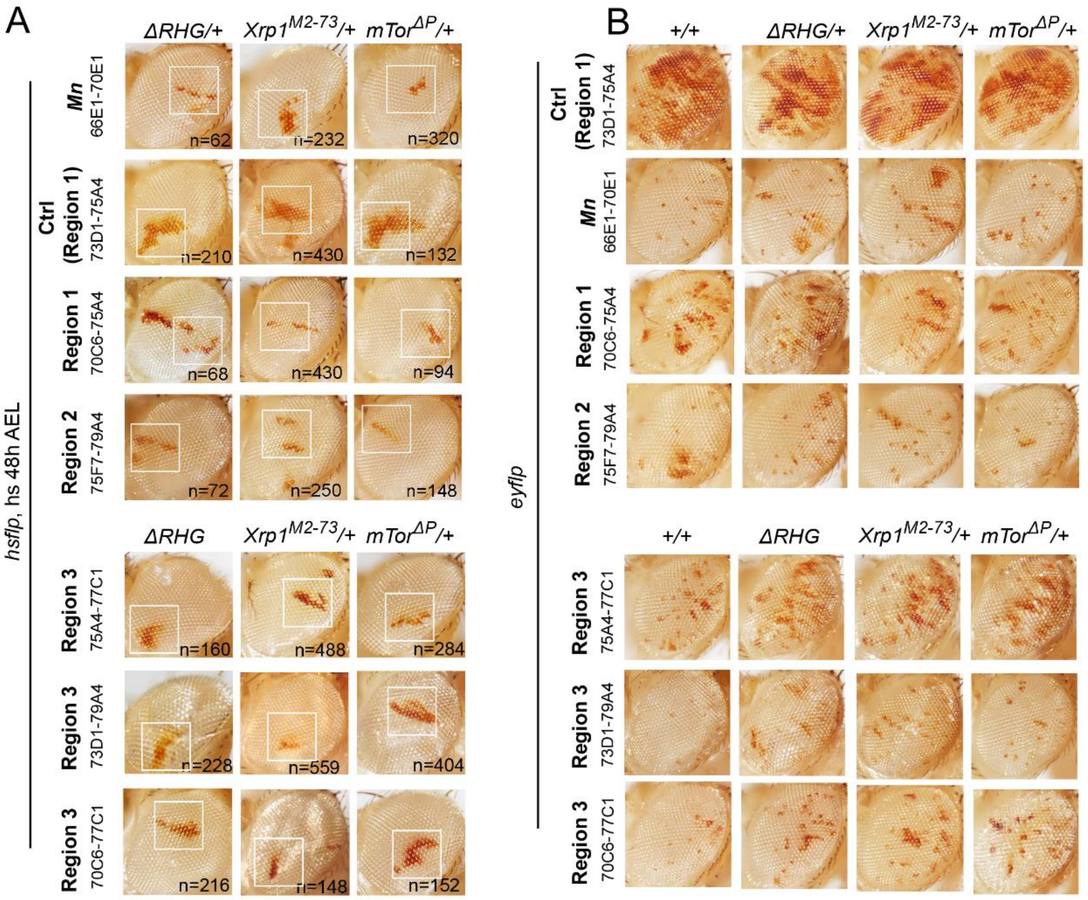


**Figure S3. Cells carrying segmental monosomies show signs of out-competition** (related to Figure 3)

(**A**, **B**) Adult eyes with clones of cells labeled in red, induced acutely at 48 h AEL (**A**) or chronically with *ey-FLP* (**B**) and bearing segmental monosomies spanning the indicated genomic locations. Clones were induced either in a wild-type background or in the indicated genetic backgrounds that compromise or block cell death, or half the doses of the *Xrp1* and *mTor* genes. High magnifications of the squared regions are shown in Figure 3A.

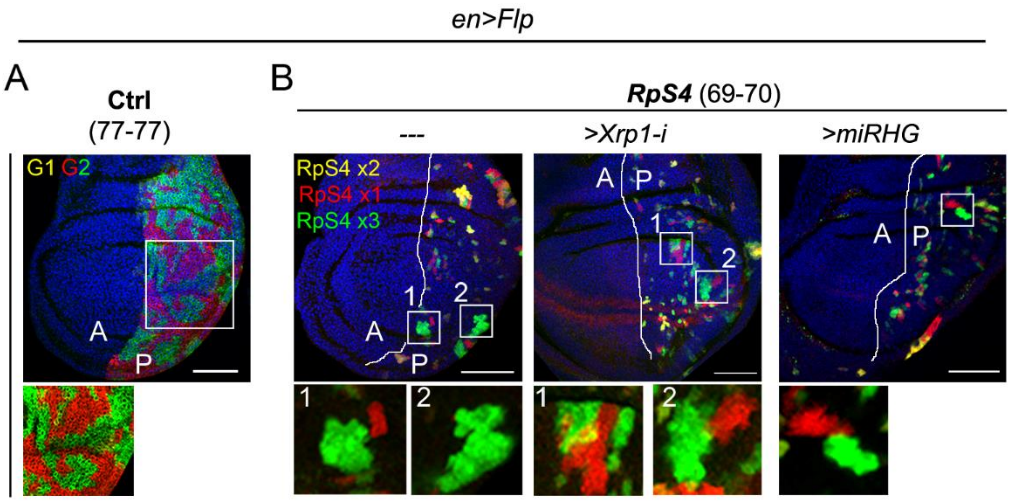


**Figure S4. The TSG technique can detect *Minute*-driven cell competition**

(**A**, **B**) Wing primordia subjected to chronic expression of *FLP* under the control of the *en-gal4* driver. Due to the high recombination efficiency of FRTs located in the same genomic location, all control clones in **A** were either red or green due to the G1 products of recombination marked in yellow resolving into segmental monosomies and trisomies by further FRT-driven recombination in G2. In **B**, clones with one (red), two (yellow) and three (green) doses of the *RpS4* gene are observed. As indicated, tissues expressed an RNAi form of *Xrp1* or the *miRHG* transgene in the posterior (P) compartment. High magnification of the squared regions is shown in the lower panels. Scale bars, 50 µm.

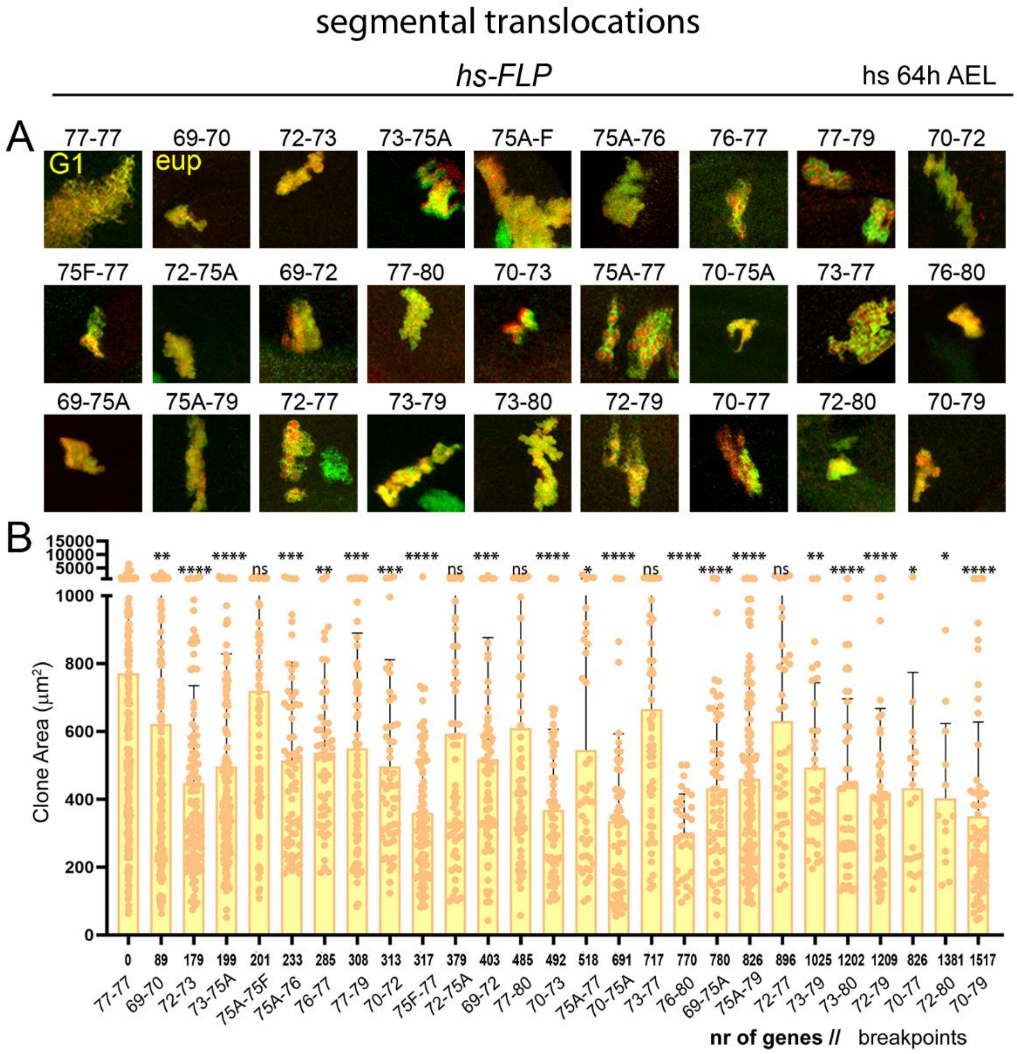


**Figure S5. Effects of segmental translocation on clonal growth** (related to Figure 5)

(**A**) Representative examples of clones of cells in wing primordia bearing segmental translocations spanning the indicated genomic locations and induced at the indicated developmental time. (**B**) Plot representing the impact of the size (in number of protein-encoding genes) of the segmental translocation on clone size (in $\mu m^2$). Genomic breakpoints of these translocations are indicated. Mean and SD are shown. Two-way ANOVA with Šidák correction for multiple comparisons test was performed. ns, not significant (p>0.05); * p ≤ 0.05; ** p ≤ 0.01; *** p≤ 0.001; and **** p ≤ 0.0001.

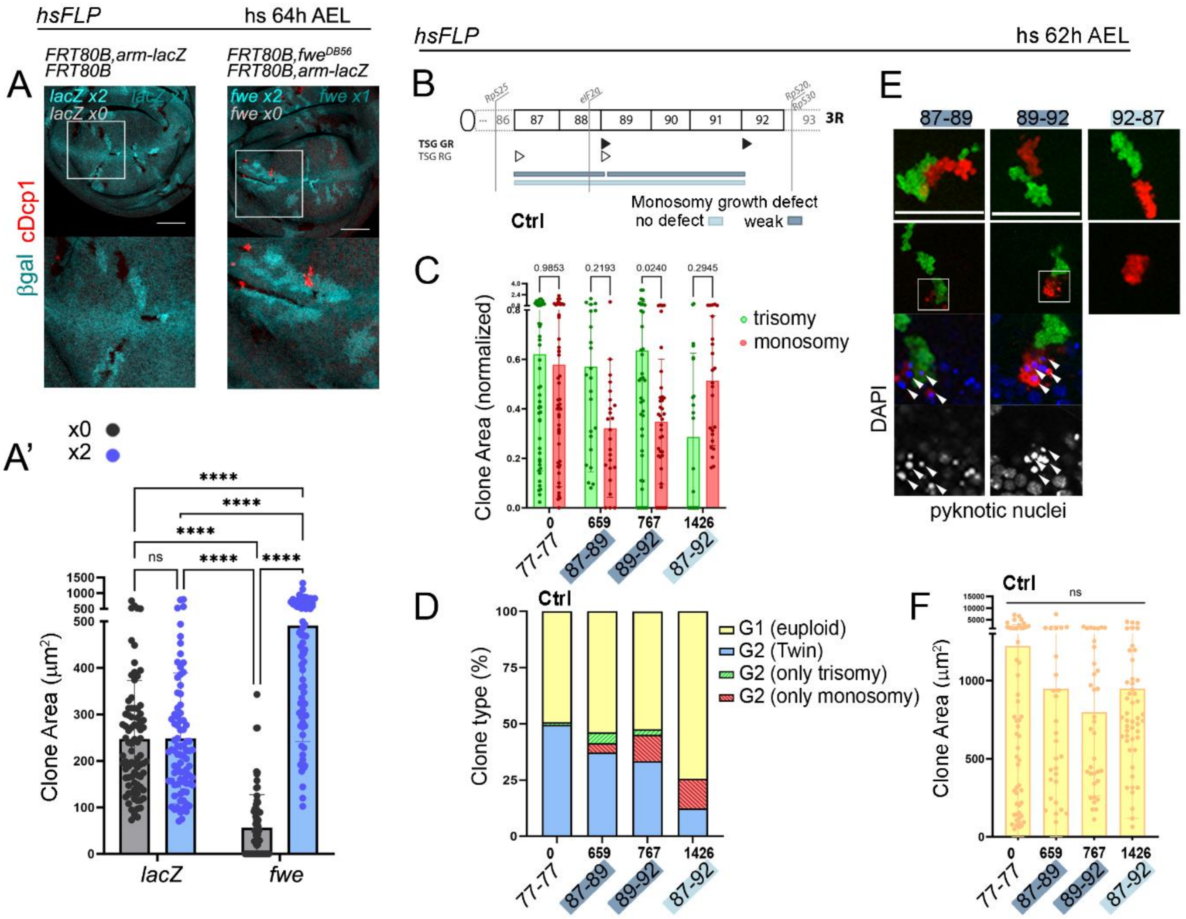


**Figure S6. Cell interactions shape the behavior of aneuploid cells: cell competition and compensation** (related to Figure 6)

(**A**) Wing disc with clones of wild type cells (left) or cells mutant for the *fwe* gene (right) marked by the absence of β-gal expression (cyan) and labeled to visualize cleaved Dcp1 (cDCp1, red). (**B**) Drawing of the right arm of the third chromosome and genomic coverage of segmental monosomies and trisomies produced. Genomic location of Ribosomal Protein (Rp)-encoding genes and *eIF2γ*, and the two types of TSG-FRTs (TSG-GR, black triangles, and TSG-RG, white triangles) used to simultaneously induce monosomic and trisomic cells are shown. Color code corresponds to the effect on growth of segmental monosomies. (**A'**, **C**, **D**, **F**) Plots representing area (in $\mu m^2$, **A'**, **F**, or normalized to the one of euploid cells, **C**) of control clones (grey in **A'**, first data set in **C**), clones with different copies of the fwe gene (**A'**), and clones bearing monosomies (red), trisomies (green) or translocations (yellow) spanning the indicated genomic locations and induced at the indicated developmental times (**C**), and distribution of the indicated types of clones (**D**). Mean and SD (A', **C**, **F**) and average (**D**) are shown. Two-way ANOVA with Šidák correction for multiple comparisons test was performed in **A'**, **C**, **F**. ns, not significant ($p>0.05$); * $p \leq 0.05$; ** $p \leq 0.01$; *** $p \leq 0.001$; and **** $p \leq 0.0001$. (**E**) Representative examples of clones of cells in wing primordia bearing segmental monosomies (red) or trisomies (green) spanning the indicated genomic locations and induced at the indicated developmental times. Arrowheads point to pyknotic nuclei labeled by DAPI (blue or white). Scale bars in **A**, **E**, 50 µm.

| FRT type | Line ID | Origin | insertion | orientation respect to chr |
|---|---|---|---|---|
| Rs5(+) | 126198 | DGRC | 70E1 | - |
| Rs5(+) | 126092 | DGRC | 72B2 | - |
| Rs5(+) | 125905 | DGRC | 73B1 | - |
| Rs5(+) | 126251 | DGRC | 73E5 | - |
| Rs5(+) | 126206 | DGRC | 75A4 | - |
| Rs5(+) | 125973 | DGRC | 77C1 | - |
| Rs5(+) | 125605 | DGRC | 78F3 | - |
| Rs5(+) | 125886 | DGRC | 78D5 | - |
| Rs5(+) | 126103 | DGRC | 79A2 | - |
| Rs5(+) | 126201 | DGRC | 79A4 | - |
| Rs5(+) | 123975 | DGRC | 80C2 | - |
| Rs3(-) | 124054 | DGRC | 70C6 | - |
| Rs3(-) | 123520 | DGRC | 70E5 | - |
| Rs3(-) | 123418 | DGRC | 71B1 | - |
| Rs3(-) | 123026 | DGRC | 71E1 | - |
| Rs3(-) | 123095 | DGRC | 73D1 | - |
| Rs3(-) | 124049 | DGRC | 75F7 | - |
| Rs3(-) | 124213 | DGRC | 77E4 | - |
| Rs3(-) | 123708 | DGRC | 78C9 | - |
| Rs3(-) | 124172 | DGRC | 78F3 | - |
| Rs3(-) | 124151 | DGRC | 79A4 | - |

**Table S1. List of RS (Rearrangement Screening) FRTs used in this work** (related to Figure 1)

RS-FRTs (RS3(-) or RS5(+)) used in this work were located in intergenic regions in the indicated genomic regions. Line ID and origin (DGRC, Drosophila Genomics Resource Center) is also indicated.

| Genotype | | Label | Pos RS3 | Pos RS5 | RS3 DGRC ID | RS5 DGRC ID | RS3 nt | RS5 nt | Diff(bp) | Mb | N genes |
|---|---|---|---|---|---|---|---|---|---|---|---|
| *eyflp; 73-75* | *hsflp; 73-75* | **73-75** | 73D1 | 75A4 | 123095 | 126206 | 16800857 | 17850544 | 1049687 | 1,049687 | 166 |
| *eyflp; 70-72B2* | *hsflp; 70-72B2* | **70-72B2** | 70E5 | 72B2 | 123520 | 126092 | 14625700 | 15948261 | 1322561 | 1,322561 | 204 |
| *eyflp; 71-73B1* | *hsflp; 71-73B1* | **71-73B1** | 71E1 | 73B1 | 123026 | 125905 | 15525670 | 16605347 | 1079677 | 1,079677 | 247 |
| *eyflp; 77-78* | *hsflp; 77-78* | **77-78** | 77E4 | 78F3 | 124213 | 125605 | 20723348 | 21815069 | 1091721 | 1,091721 | 247 |
| *eyflp; 70E5-72D9* | *hsflp; 70E5-72D9* | **70E5-72D9** | 70E5 | 72D9 | 123520 | 125972 | 14625700 | 16157381 | 1531681 | 1,531681 | 256 |
| *eyflp; 77-79* | *hsflp; 77-79* | **77-79** | 77E4 | 79A4 | 124213 | 126201 | 20723348 | 21935345 | 1211997 | 1,211997 | 266 |
| *eyflp; 75-77* | *hsflp; 75-77* | **75-77** | 75F7 | 77C1 | 124049 | 125973 | 19094051 | 20394712 | 1300661 | 1,300661 | 272 |
| *eyflp; 70C6-72B2* | *hsflp; 70C6-72B2* | **70C6-72B2** | 70C6 | 72B2 | 124054 | 126092 | 13932268 | 15948261 | 2015993 | 2,015993 | 282 |
| *eyflp; 71-73E5* | *hsflp; 71-73E5* | **71-73E5** | 71E1 | 73E5 | 123026 | 126251 | 15525670 | 17042518 | 1516848 | 1,516848 | 324 |
| *eyflp; 75A4-77* | *hsflp; 75A4-77* | **75A4-77** | 75A4 | 77C1 | 124168 | 125973 | 17850477 | 20394712 | 2544235 | 2,544235 | 468 |
| *eyflp; 70-73* | *hsflp; 70-73* | **70-73** | 70C6 | 73E5 | 124054 | 126251 | 13932268 | 17042518 | 3110250 | 3,11025 | 537 |
| *eyflp; 75-79* | *hsflp; 75-79* | **75-79** | 75F7 | 79A4 | 124049 | 126201 | 19094051 | 21935345 | 2841294 | 2,841294 | 608 |
| *eyflp; 66-70* | *hsflp; 66-70* | **66-70** | 66E1 | 70E1 | 123215 | 126198 | 8820579 | 14530694 | 5710115 | 5,710115 | 619 |
| *eyflp; 70-75* | *hsflp; 70-75* | **70-75** | 70C6 | 75A4 | 124054 | 126206 | 13932268 | 17850544 | 3918276 | 3,918276 | 663 |
| *eyflp; 75A4-78D5* | *hsflp; 75A4-78D5* | **75A4-78D5** | 75A4 | 78D5 | 124168 | 125886 | 17850477 | 21526856 | 3676379 | 3,676379 | 708 |
| *eyflp; 75A4-78F3* | *hsflp; 75A4-78F3* | **75A4-78F3** | 75A4 | 78F3 | 124168 | 125605 | 17850477 | 21815069 | 3964592 | 3,964592 | 785 |
| *eyflp; 73-79* | *hsflp; 73-79* | **73-79** | 73D1 | 79A4 | 123095 | 126201 | 16800857 | 21935345 | 5134488 | 5,134488 | 970 |
| *eyflp; 71-77* | *hsflp; 71-77* | **71-77** | 71B1 | 77C1 | 123418 | 125973 | 15007510 | 20394712 | 5387202 | 5,387202 | 999 |
| *eyflp; 70-77* | *hsflp; 70-77* | **70-77** | 70C6 | 77C1 | 124054 | 125973 | 13932268 | 20394712 | 6462444 | 6,462444 | 1131 |
| *eyflp; 70-78* | *hsflp; 70-78* | **70-78** | 70C6 | 78D5 | 124054 | 125886 | 13932268 | 21526856 | 7594588 | 7,594588 | 1371 |
| *eyflp; 70-79* | *hsflp; 70-79* | **70-79** | 70C6 | 79A4 | 124054 | 126201 | 13932268 | 21935345 | 8003077 | 8,003077 | 1467 |
| *eyflp; 70-80* | *hsflp; 70-80* | **70-80** | 70C6 | 80C2 | 124054 | 123975 | 13932268 | 22991401 | 9059133 | 9,059133 | 1725 |

**Table S2. List of RS (Rearrangement Screening) FRT combinations in cis used in this work** (related to Figure 2)

List of recombinant fly lines bearing 21 different combinations of RS5r and RS3r-FRTs located in the same chromosome (*in cis*), where RS5r is distal to RS3r on the 3L (*RS5r RS3r*). Source of FLP expression, genomic location of RS-FRTs (cytogenetic and molecular), distance between RS-FRTs (in bp, Mb, and number of genes) are indicated.

| FRT type | Line ID | Origin | insertion | orientation respect to chr |
|---|---|---|---|---|
| TSG GR | 43615 (MiMIC ) | BSDC | 69F1 | + |
| TSG RG | 36936 (MiMIC ) | BSDC | 70A8 | + |
| TSG GR | 36936 (MiMIC ) | BSDC | 70A8 | + |
| TSG RG | 40163 (MiMIC ) | BSDC | 72A1 | + |
| TSG RG | 36406 (MiMIC ) | BSDC | 73A5 | + |
| TSG GR | 36406 (MiMIC ) | BSDC | 73A5 | + |
| TSG RG | 56571 (MiMIC ) | BSDC | 75A1 | + |
| TSG GR | 56571 (MiMIC ) | BSDC | 75A1 | + |
| TSG GR | 43044 (MiMIC ) | BSDC | 75F1 | + |
| TSG RG | 35938 (MiMIC ) | BSDC | 76A3 | + |
| TSG RG | Griffin et al 2009 | Perrimon lab | 77C4 | + |
| TSG GR | Griffin et al 2009 | Perrimon lab | 77C4 | + |
| TSG GR | 31404 (MiMIC ) | BSDC | 79A4 | + |
| TSG GR | 58655 (MiMIC ) | BSDC | 80B1 | + |
| TSG GR | 53833 (MiMIC ) | BSDC | 87A4 | + |
| TSG RG | 53833 (MiMIC ) | BSDC | 87A4 | + |
| TSG GR | 44927 (MiMIC ) | BSDC | 89A1 | + |
| TSG GR | 44869 (MiMIC ) | BSDC | 92F6 | + |

**Table S3. List of TSG (Twin Spot Generator) FRTs used in this work** (related to Figure 4)

TSG-FRTs (RG or GR) were injected into the indicated MiMIC lines. Line ID and origin (BSDC, Bloomington Drosophila Stock Center), genomic location and orientation of the insertion with respect to the chromosome are also indicated.

**Table S4. List of genes included in the segmental aneuploidies** (related to Figure 4-7)

Genes included in each of the segmental aneuploidies generated by the indicated RS-FRTs and TSG-FRTs. Genomic region (1, 2 or 3, first column), observed cellular behavior (second column), RS-FRT ID and TSG-FRT genomic location (third to sixth columns), gene and transcript ID, gene start and end, gene description, karyotype band and gene name (seventh to thirteenth columns) are indicated.

**Table S5. File containing the parameters that have been quantified and the statistical details** (related to Figures 1-7 and S1, S2, S3, S5, S6).

Quantification and statistical analysis of recombination efficiency, and growth and survival parameters of clones of cells bearing segmental aneuploidies.
